# Massively parallel characterization of regulatory dynamics during neural induction

**DOI:** 10.1101/370452

**Authors:** Fumitaka Inoue, Anat Kreimer, Tal Ashuach, Nadav Ahituv, Nir Yosef

## Abstract

The molecular components governing neural induction remain largely unknown. Here, we applied a suite of genomic and computational tools to comprehensively identify these components. We performed RNA-seq, ChIP-seq (H3K27ac, H3K27me3) and ATAC-seq on human embryonic stem cells (hESCs) at seven early neural differentiation time points (0-72 hours) and identified thousands of induced genes and regulatory regions. We analyzed the function of ~2,500 selected regions using massively parallel reporter assays at all time points. We found numerous temporal enhancers that correlated with similarly timed epigenetic marks and gene expression. Development of a prioritization method that incorporated all genomic data identified key transcription factors (TFs) involved in neural induction. Individual overexpression of eleven TFs and several combinations in hESCs found novel neural induction regulators. Combined, our results provide a comprehensive map of genes and functional regulatory elements involved in neural induction and identify master regulator TFs that are instrumental for this process.

**One Sentence Summary:** Using numerous genomic assays and computational tools we characterized the dynamic changes that take place during neural induction.

## Main Text

Pluripotent cells differentiate into a neural lineage as a default when BMP and TGFβ signals are inhibited (*1*). Neural induction is the initial step of this process, priming these cells to later become neural progenitor cells (NPCs). Human embryonic stem cells (hESCs) can also be differentiated into NPCs when BMP and TGFβ inhibitors are added into culture media (*2*). As such, neural induction of hESC is widely used as a model system to study the mechanism of neural differentiation and neurodevelopmental diseases. However, we currently have a limited understanding of the molecular components governing this process; in particular, the regulatory elements orchestrating it. Changes in enhancer activity are thought to play a pivotal role in cell fate specification (*3*). For example, multiple temporal enhancers are thought to sequentially function in orthodenticle homeobox 2 (*Otx2*) gene expression in epiblast, anterior neuroectoderm and fore-and midbrain during development (*4-6*).

Mutations in genes and regulatory elements involved in neural induction and development have been associated with human disease. For example, dysfunction of cortical GABA neurons in schizophrenia begins during prenatal development (*7*). Similarly, autism spectrum disorders (ASD) are associated with *de novo* mutations in developmental genes (*8*) and alterations in canonical Wnt signaling in developing embryos (*9*). Furthermore, copy number variations (CNVs) overlapping neurodevelopmental genes linked to ASD and schizophrenia were found to be associated with attention-deficit hyperactivity disorder (ADHD) (*10*) and genetic factors that lead to prefrontal-subcortical network dysfunction are thought to be associated with pediatric bipolar disorder (*11*). In addition, the majority of disease-risk loci discovered through genome-wide association studies (GWAS) in general and specifically for neuropsychiatric and neurodevelopmental disorders reside in noncoding regions (*12-14*), suggesting an important role for enhancers in disease susceptibility. Several large-scale mapping efforts have characterized in a genome-wide manner the transcriptional and epigenetic landscape of hESC-derived NPCs or neural tissues and have annotated numerous genes and potential regulatory elements that could be important in neural differentiation (*15-21*). However, while these studies have identified putative regulatory elements, they have not comprehensively analyzed them for their function. Furthermore, none of these genomic studies focused on the early stages of neural differentiation when neural induction takes place. Thus, the intrinsic mechanism that governs neural induction remains largely unknown.

Here, we set out to generate a genomic map of the transcriptional and epigenetic landscape of neural induction, and then couple these observations with comprehensive functional assays, with the aim of identifying the key regulatory elements (enhancers, transcription factors) involved in this process. To this end, we measured gene expression levels (RNA-seq), chromatin accessibility (ATAC-seq), and histone modifications (ChIP-seq) indicative of active (H3K27ac) and repressed (H3K27me3) regions at seven time points, spanning the early stages of neural differentiation (0-72 hours). We first used these datasets to identify genes that are regulated during (and thus potentially associated with) neural induction, as well as their associated regulatory regions. Next, we prioritized and tested 2,464 candidate regions using a lentivirus-based massively parallel reporter assay (lentiMPRA;(*22*)) for regulatory activity. LentiMPRA found 62% (1, 547/2,464) of the assayed sequences to be temporally active, namely, capable of driving reporter transcription while showing differential activity over time. The majority (92%) of these sequences are novel enhancers that were not tested for enhancer activity in previous studies. Importantly, the activity of over two thirds of the temporal enhancers matched with the temporal profiles observed in the endogenous genome, either in terms of chromatin accessibility, H3K27 acetylation or mRNA expression. We integrated all of the resulting data modalities (genomics maps and MPRA) to computationally infer the activity of transcription factors (TFs) over time and nominate candidate TFs that could be important drivers of neural induction. Overexpression of eleven of these TFs along with selected combinations, found three novel TFs (OTX2, LHX5 and IRX3) that can induce neural differentiation in hESCs. Combined, our work provides a comprehensive gene and regulatory map of sequences involved in neural induction and identifies novel functional enhancers and TFs involved in this process.

## The neural induction-associated transcriptome

To characterize the neural induction transcriptome, we performed deep RNA sequencing (average of 200 million reads per replicate) on undifferentiated H1-ESCs (0 hour) and six different time points of early neural differentiation (3, 6, 12, 24, 48, and 72 hours) following dual-Smad inhibition (*2*). As expected, we observed neural marker genes (*SOX1*, *PAX6*, *OTX2*, *LHX5*, *IRX3*, *POU3F2*, *DLK1*, *MAP2* and *CDH2*) to be upregulated after 12 hours (**Fig. 1A**), with limited expression changes in mesendoderm (*EOMES*), mesoderm (*T* and *TBX6*), endoderm (*SOX17* and *GATA4*), and neural crest markers (*FOXD3* and *SNAI1/2*). Pluripotent markers (*NANOG*, *POU5F1*) and direct targets of TGFβ and BMP signaling (*SMAD7*, *ID1*, *LEFTY2*) were downregulated and immediate early genes (*ATF3*, *FOS, FOSB* and *EGR1/2/3*) were transiently upregulated at 3 hours, corresponding to the cell’s stress response against differentiation stimuli. For a more general analysis, we used two different methods for the identification of differential expression over time (SigmoDE (*23*) and DESeq2 with time covariates (*24*)) and considered only genes that were called by both [using a false discovery rate (FDR) cutoff of 1% and 5% respectively; **Methods**]. Altogether, we found 2,172 genes that are differentially expressed over time (henceforth referred to as *temporal genes*), with 85% of these genes being induced at some point in time (**Fig. 1B**). Gene set enrichment analysis (*25*) of the resulting temporal clusters found that genes induced at the early time points (0-12 hours, FDR<0.05 hypergeometric test) are enriched for regulation of multicellular organismal development, indicating that pluripotent response genes are enriched in these processes. Conversely, genes induced at later time points (>24 hours, FDR<0.05) are enriched for neurogenesis processes, consistent with the progression of the cells toward a neural lineage fate (**table S1**). Combined, our transcriptomic analyses validated the ability of the dual-Smad inhibition protocol to obtain the expected neural trajectory and provides a catalog of genes involved in neural induction.

**Fig. 1.**
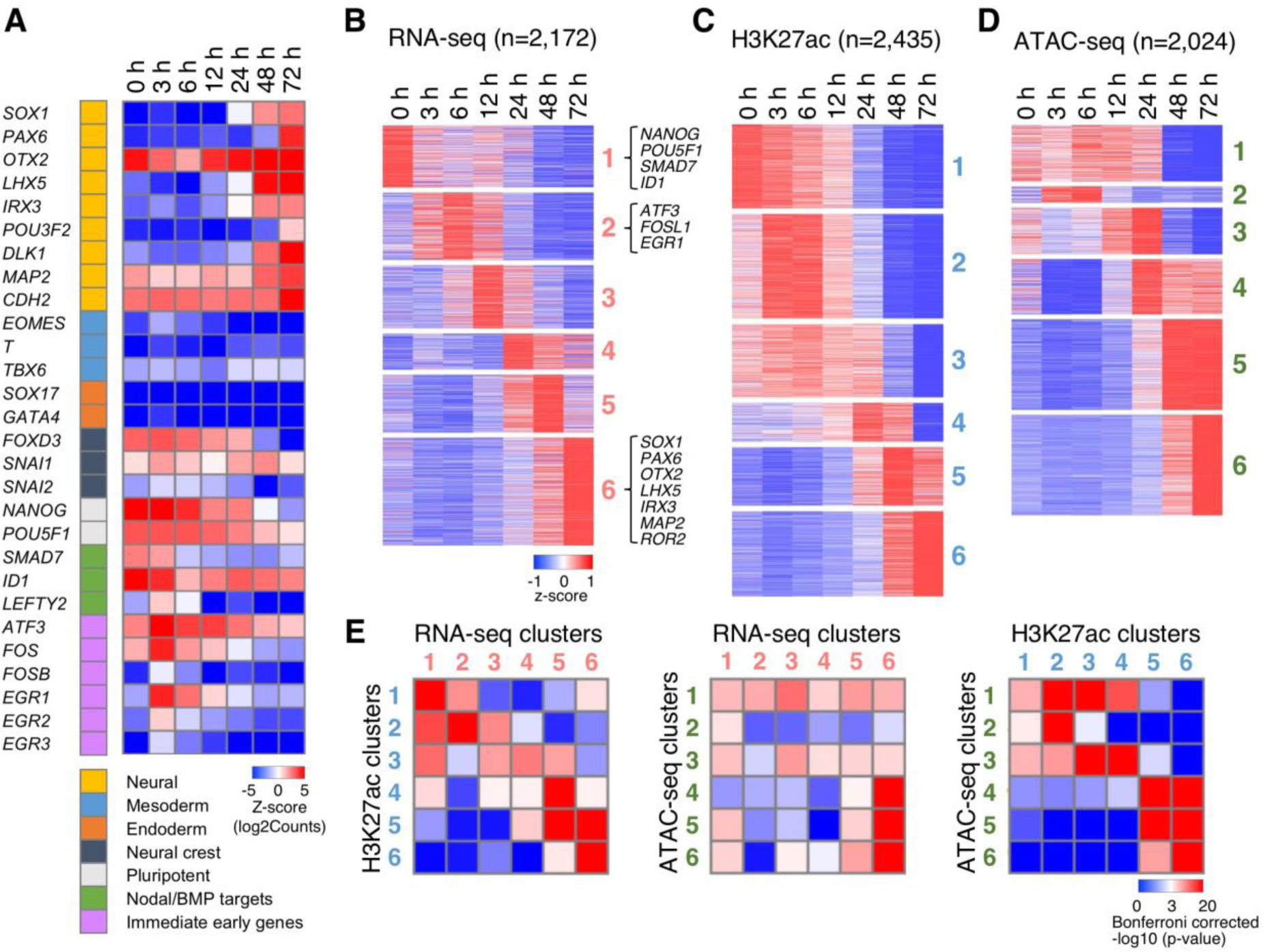
The dynamic changes of ATAC-seq, ChIP-seq and RNA-seq peaks are sequentially correlated. (**A**) Scaled read count (log2, averaging over three biological replicates) per time point of genes from seven representative groups (Neural, Mesoderm, Endoderm, Neural crest, Pluripotent, Nodal/BMP targets and immediate early genes). (**B-D**) Heat map of scaled read counts (log2, averaged over three biological replicates and standardized per row) of temporal genes and genomic regions, showing data from RNA-seq (**B**), H3K27ac ChIP-seq (**C**) and ATAC-seq (**D**). The loci in each assay were clustered into six groups based on their temporal patterns. (**E**) Overlap between the temporal clusters in the three data modalities. Shown are Bonferroni corrected p-values of a hypergeometric test. The overlap is computed either at the region level (ATAC-seq vs. ChIP-seq) or at the gene level (ATAC/ChIP-seq vs. RNA-seq; regions in the former assays are represented by their nearest gene).

## The neural induction-associated regulome

To comprehensively identify candidate enhancers involved in regulating these genes (and thus driving neural induction), we performed ATAC-seq as well as ChIP-seq for the active histone mark H3K27ac and the silencing mark H3K27me3 at all seven time points. We then identified regions that are enriched (i.e., peak regions) in each one of these assays (FDR< 0.05) by analyzing each time point separately and then taking the merged set of peaks over all time points. Overall, we identified 40,486 ATAC-seq peaks, 40,170 H3K27ac peaks and 4,446 H3K27me3 peaks that are enriched in at least one time point. To exclude potentially inactive regions from further analysis, we removed H3K27ac peaks that overlap a H3K27me3 peak at all the time points in which that peak was detected. This resulted in a filtered set of 40,042 H3K27ac peaks, indicating that the two chromatin marks have little overlap in our data. Conversely, we observed a substantial overlap between the H3K27ac peaks and the ATAC-seq peaks, with an overall 60% (23,294) of the H3K27ac peaks overlapping an accessible region at the same point in time. Using strict criteria [FDR< 0.05; FDR<0.01 **Methods; table S1**; (*23, 24*)], as in the gene expression analysis, we found 2,435 ATAC-seq and 2,024 H3K27ac peaks that were differentially enriched between time points, henceforth referred to as *temporal H3K27ac* or *ATAC-seq peaks*. Similar clustering of H3K27me3 peaks showed weaker temporal signal (**Methods**) and a smaller number of temporal peaks (**fig. S1A**).

We next set out to study the association between the temporal changes observed at the epigenome level, and those observed at the gene expression level. We clustered the two sets of temporal regions (in terms of accessibility and H3K27ac) into several prototypical patterns (**Fig. 1C, D**) as we have done for the temporal genes **(Fig. 1B).** Functional enrichment analysis using the Genomic Regions Enrichment of Annotations Tool [GREAT; (*26*) FDR<0.05] on the accessibility and H3K27ac clusters was overall consistent with the results observed with the gene expression clusters, with an enrichment for pluripotent factors and nervous system development processes in early and late response regions respectively (**table S1**). Interestingly, we observed that the temporal changes to the epigenome were highly correlated to each other (**Fig. 1E and fig. S1B**). Furthermore, for a large fraction of the induced genes, chromatin accessibility was found to be acquired first or simultaneously with H3K27ac modification followed by an increase in mRNA expression (using the expression of the nearest gene; **Fig. 1E**). For example, the DNA accessibility cluster 4 that peaks at 24 hours showed the strongest overlap with H3K27ac clusters 5-6 that peak at 48-72 hours and this cluster significantly overlaps (in terms of genes; p-value<0.0014; hypergeometric test) with gene expression cluster 6 which peaks at 72 hours (**Fig. 1E**). Specifically, examination of potential enhancers within these clusters that are located near microtubule associated protein 2 (*MAP2*), a gene that is involved in microtubule assembly (*27*), or receptor tyrosine kinase like orphan receptor 2 (*ROR2*), that regulates the maintenance of NPCs (*28*), found them to be enriched for ATAC-seq signal at 12-24 hours, H3K27ac signal at 48-72 hours and their expression to peak at 72 hours (**Fig. S2**). Combined, these results suggest that regions that are associated with changes to chromatin structure during neural induction are statistically related to changes in gene expression.

## Neurological disorder associated variants are enriched in temporal H3K27ac marked sequences

As genes and regulatory elements involved in neural development may be associated with neurological disorders, we tested whether our neural induction regulome overlaps disease-associated variants. We first analyzed whether our temporal accessibility or H3K27ac peaks are specifically enriched for GWAS variants associated with neurological disorders using the complete set of peaks (temporal and non-temporal) as background and variants associated with height as negative controls. We observed a significant enrichment for H3K27ac (but not accessibility) temporal peaks with neurological disorders (**table S2**; **Methods**; p-value<0.05; Fisher’s exact test) but not with height variants. Specifically, we observe significant enrichment when examining variants associated with a combined set of neuropsychological disorders (schizophrenia, attention-deficit/hyperactivity disorder, autism spectrum disorder, bipolar disorder and major depressive disorder) as well as enrichment when examining for individual disorders (i.e. Bipolar and Psychosis disorders). As the smaller size of ATAC-seq peaks might account for the lack of enrichment in ATAC-seq temporal peaks, we expanded the ATAC-seq peaks to the average size of H3K27ac peaks, but observed similar results.

Expression quantitative trait loci (eQTLs) mark variants that can be associated with modulating the regulation of nearby genes. We tested for overlap between eQTLs found in various tissues (*29*) and our temporal ATAC-seq or H3K27ac peaks. We found the temporal H3K27ac peaks to be significantly enriched for eQTL variants (*30*) in general and specifically for those from brain tissues (*29*) (**table S2**; **Methods**; p-value<0.05; Fisher’s exact test using all peaks, temporal and non-temporal, as background). Similarly to GWAS variants, we did not observe an enrichment of eQTLs in temporal ATAC-seq peaks even upon their expansion. Combined, these results suggest that our temporal H3K27ac regions could be functional enhancers that harbor neurological disease risk variants. They also suggest that temporal changes to the chromatin early in the differentiation process can be used as a tool for identifying potentially functional regions (more so than a single time point).

## LentiMPRA identifies regulatory regions that are active during neural induction

In order to test whether our candidate regulatory sequences can in fact induce temporal transcriptional response, we carried out lentiMPRA at all seven time points. Overall, we investigated 2,464 candidate sequences, covering both promoters (N=386 (15.7%)) and putative enhancers (N=2,078 (84.3%)). As the number of potential candidate regulatory regions is large, we developed a prioritization scheme to select the set of assayed regions, (**Fig. 2A; table S3; Methods**) using the following criteria: 1) Manually curated list of enhancers that are next to genes involved in neural differentiation (N=102; **table S3**); 2) Sequences that overlap a temporal H3K27ac ChIP-seq peak that also overlap an ATAC-seq peak (not necessarily temporal) and that their closest gene shows increased expression due to neural induction (N=1,596); 3) Sequences that overlap non-temporal H3K27ac peaks and temporal ATAC-seq peaks and their closest gene shows increased expression due to neural induction (N=441); 4) Among the regions not included in the first three groups, we select sequences that showed the strongest difference in signal of either H3K27ac ChIP-seq, ATAC-seq or mRNA of the closest genes (N=132; comparing either 0 vs. 3 hours or 0 vs. 72 hours); and 5) Positive control sequences (N=193) that included previously reported sequences that were validated forebrain enhancers in the VISTA Enhancer Browser (Visel et al., 2009) (N=105), sequences near pluripotent factors (N=42) and commonly used positive controls from the ENCODE project (N=46) (**table S3**). Notably, the criteria were applied sequentially (in the order in which they were described), and the respective sets of candidate enhancers are mutually exclusive. For negative controls, we randomly selected 200 of our candidate sequences and shuffled their nucleotides obtaining scrambled sequences. Overall, we chose 2,664 sequences using this process. As our assayed sequences were 171 base pairs (bp) long, due to oligonucleotide synthesis limitations, we chose the 171bp window within a peak of interest by maximizing the number of motifs in it (**Methods**; (*31, 32*)).

**Fig. 2.**
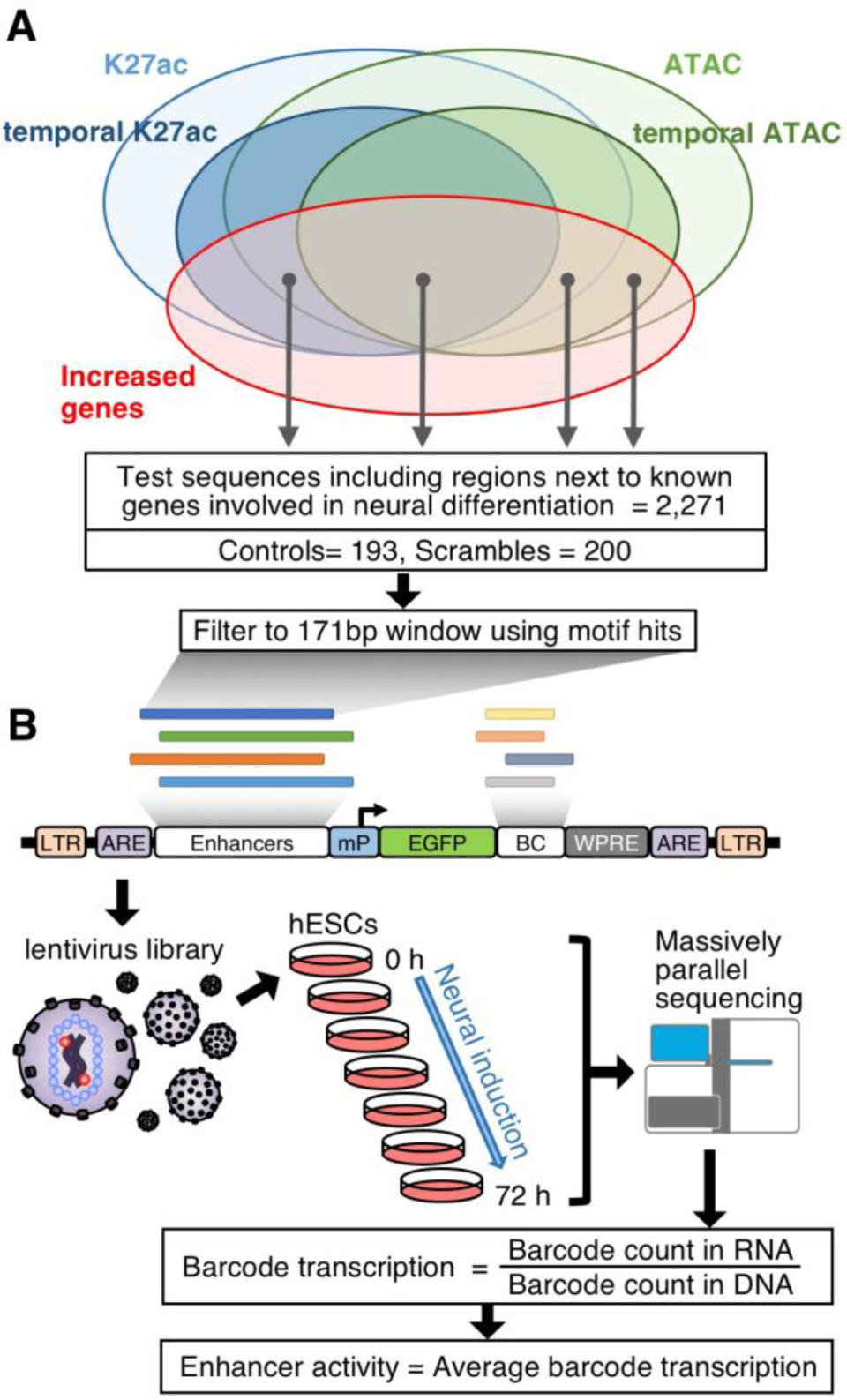
Experimental design of lentiMPRA. (**A**) Sequence selection for lentiMPRA. 2,271 test regions were selected from the following four groups: 1) regions next to known genes involved in neural differentiation; 2) regions that overlap some ATAC-seq peak and a temporal H3K27ac peak and reside near genes that show increased expression due to neural induction; 3) regions that overlap a temporal ATAC-seq peak and a non-temporal H3K27ac peak and reside near genes that show increased expression due to neural induction; 4) regions with strong differential enrichment for either H3K27ac ChIP-seq, ATAC-seq or near genes that showed strong differential expression in our studies that do not overlap with the first three groups. 193 positive control regions and 200 scramble sequences (negative controls) were also included. (**B**) Schematic showing lentiMPRA design. Putative enhancer sequences along with a 15 bp barcodes were synthesized on a custom array, and cloned into a lentiMPRA vector. To ensure robustness we designed for each enhancer ninety different vectors, each with a different barcode. The library was packaged into lentivirus and infected into hESCs. The infected cells were cultured for 3 days, to allow genomic integration. DNA and nuclear RNA were extracted at seven time points (0, 3, 6, 12, 24, 48, 72 h) and subjected to sequencing followed by estimation of transcriptional activity.

The selected oligonucleotides were generated and cloned upstream of a minimal promoter (mP) and EGFP reporter gene into a lentivirus-based enhancer assay vector (**Fig. 2B**) as previously described (*22*). Previous work has shown this assay to also provide a good indication for promoter activity (*33, 34*). Each individual enhancer sequence was designed to be associated with 90 different 15-bp barcodes, thus allowing robust evaluation of the pertaining expression output. In total 239,760 sequences (2,664 enhancers x 90 barcodes) were included in the library (**Fig. 2B**).

The cloned library was sequenced in order to evaluate the quality of the designed oligonucleotides and the representation of individual barcodes (**Methods; fig. S3**). We found small deviations of the sequence design from observed length (**fig. S3A**) and distributions of alignment errors to be minimal and evenly distributed along the sequence (**fig. S3B**). These results were comparable to a previous library generated in a similar manner (*22*).

hESCs were infected with the library with an average of 5-8 integrations per cell (**fig. S4**), cultured for 3 days to clean out for unintegrated lentivirus and then subsequently induced into a neural lineage via dual-Smad inhibition. LentiMPRA was performed at all seven time points of neural differentiation with three replicates (two biological replicates, one of which was split into two technical replicates; see **Methods, figs. S5-10** and **table S4**). Due to the short time spans between some conditions, we collected nuclear RNA in all time points so as to detect their immediate expression. We observed an average of 70 barcodes per candidate enhancer sequence in each replicate (out of 90 barcodes programmed on the array; **fig. S6**). These results were comparable to a previous library generated in a similar manner (*22*). By aggregating these barcodes (**Methods**), we were able to get highly reproducible results across both technical and biological replicates (**figs. S7; S8A-C**). We then combined replicates to produce a normalized RNA/DNA ratio for each enhancer (**Methods**; henceforth referred to as *MPRA signal*). We observed low correlation between RNA/DNA ratios and DNA counts, indicating that enhancer activity was not influenced by the number of DNA integrations (**fig. S9**). Examination of the signal observed for regions nominated by the different experimental design criteria found that temporal H3K27ac signal (criterion 2) provides an effective predictor of functional enhancer activity, while as expected, the negative controls showed the lowest activity (**Fig. 3A; fig. S8D**).

**Fig. 3.**
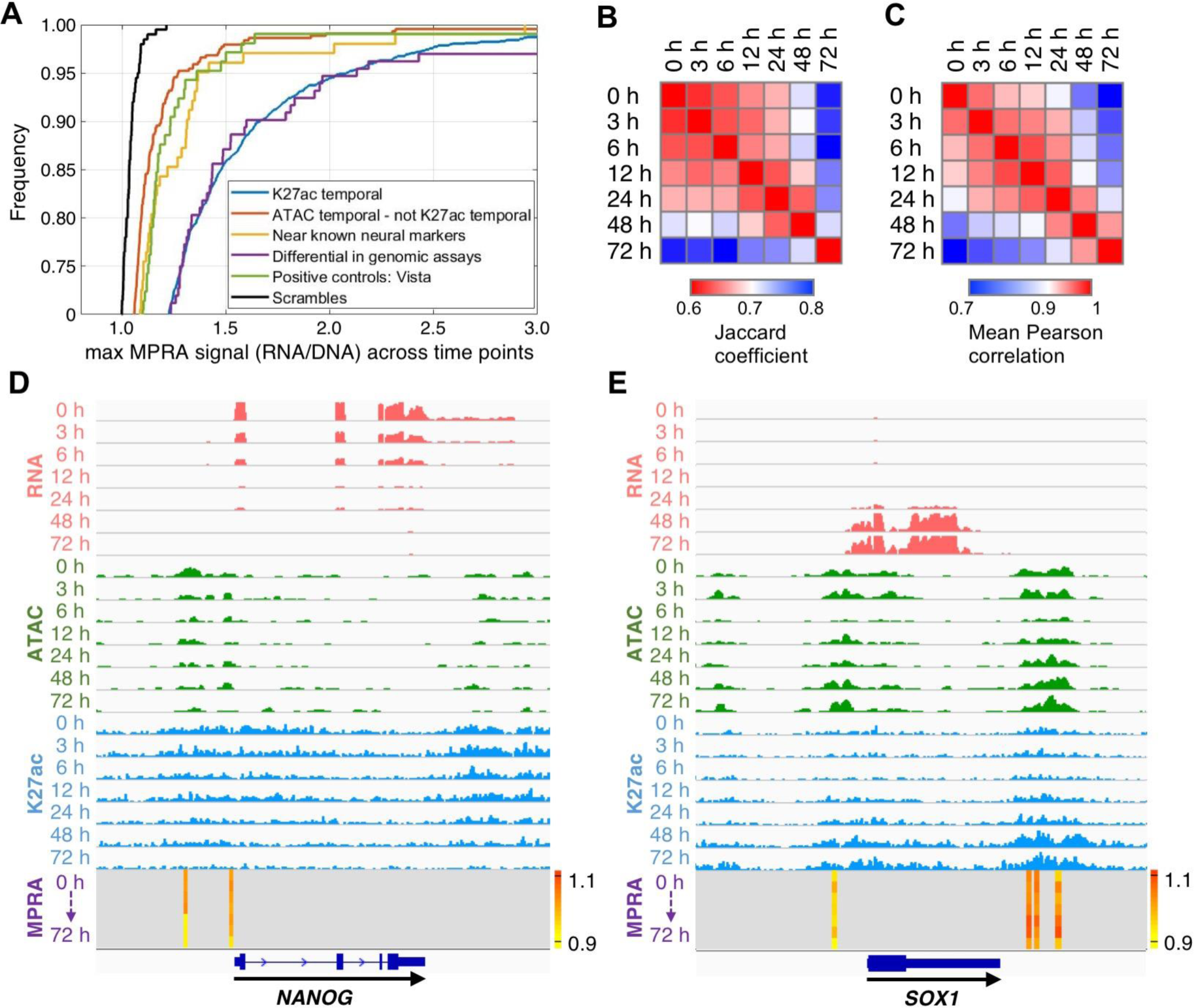
lentiMPRA signal for different enhancer types. (**A**) Cumulative distribution function indicating the frequency (y-axis) of sequences with a specific MPRA signal (x-axis; taking the maximum signal over time) revealed strong association between enhancer activity and temporal H3K27ac enrichment (design criterion 1) as well as regions that show differential activity when comparing extreme time points (0 vs. 3 hours, 0 vs. 72 hours) of H3K27ac, ATAC-seq and RNA-seq signals (design criterion 4). (**B-C**) Similarity between the MPRA signal measured at different time points, using either the intersection of the sets of significantly active regions (Jaccard coefficient; **B**), or the correlation of the signals (Pearson correlation; **C**). (**D-E**) RNA-seq (red), ATAC-seq (green), H3K27ac ChIP-seq (blue) and MPRA (RNA/DNA ratio heatmap) tracks around *NANOG* (**D**) and *SOX1* (**E**).

## LentiMPRA identifies temporal enhancers

We next set out to examine whether the enhancer activity observed in our assay changes over time, and then characterize these changes with respect to the cell-endogenous temporal processes observed in **Fig. 1**. As a starting point, we considered each time point separately and applied MPRAnalyze (**Note S1**) a new tool for statistical analysis of MPRA data developed in our group, to identify *active* enhancers, namely enhancers whose activity significantly deviates from that of the negative controls (median-based z-score; FDR < 0.05). From the 2,464 candidate sequences that we tested via lentiMPRA, 1,681 (68%) were called significant in at least one of the time points and on average 1,141 (46.3%) sequences were active per each individual time point (**table S3**).

While we saw similar levels of activity at each time point, the sets of responsive enhancers may differ greatly between time points. Reassuringly, we observed a correlation between the time points and temporal order; namely, the overlap between the sets of active enhancers (**Fig. 3B**) and the correlation of MPRA signal for all enhancers is decreased as the distance between the respective time points increases (**Fig. 3C; fig. S10**). This indicated that regulatory programs carried out by enhancers are far from fixed, but instead change over the course of neural induction. As an example, we observed that a known enhancer and the promoter of *NANOG* (*35, 36*) have activity only at the early time points (**Fig. 3D**), as expected. We also found novel enhancers near SRY-box 1 (*SOX1*) that showed increased activity at 24-48 hours (**Fig. 3E**), corresponding to the respective mRNA expression changes over time (*2*) (**Fig. 1A**).

We next carried out a more global analysis that aims to identify enhancers whose MPRA signal significantly changed over time (**Note S1**). This alternative approach pools together information from all time points rather than considering each time point individually, and therefore has the potential to identify effects that may otherwise be missed. In this analysis, the temporal activity of each candidate enhancer was compared with a null temporal behavior displayed by the set of negative controls. Regions with significantly different temporal activity were called temporally active using a likelihood ratio test (FDR< 0.05; **Methods; Note S1**). We found that 1,547 sequences out of the 2,464 we tested (63%) showed temporal enhancer activity (henceforth referred to as *temporal enhancers*). Out of these temporal enhancers 1,261 (82%) were also detected by the per-time point analysis. In the following analyses we focused on the complete set of temporal enhancers. Importantly, we observed consistent results when limiting our analyses to the smaller and more stringent set of 1,261 regions.

## Enhancer activity is consistent with the endogenous temporal profiles

We set out to test whether the MPRA signal of the temporal enhancers correlates with gene expression and endogenous enhancer marks. To this end, we clustered the temporal enhancers into four patterns of activity: 1) early (mainly active at 0-6 hours); 2) mid-early (primarily active at 12-24 hours); 3) mid-late (mainly active at 24-48 hours); and 4) late response (primarily active at 48-72 hours) (**Fig. 4A**). To facilitate direct comparison to the temporal profiles observed in the endogenous genome, we quantified for each temporal enhancer the expression of its closest gene and the epigenetic signal (accessibility, H3K27ac) in the respective endogenous position over time. We then stratified the resulting profiles into clusters, in a similar way to that of the MPRA (**Fig. 4B-D**) and tested the overlap between the resulting endogenous clusters (**Fig. 4B-D)** and the MPRA-based clusters (**Fig. 4A**).

**Fig. 4.**
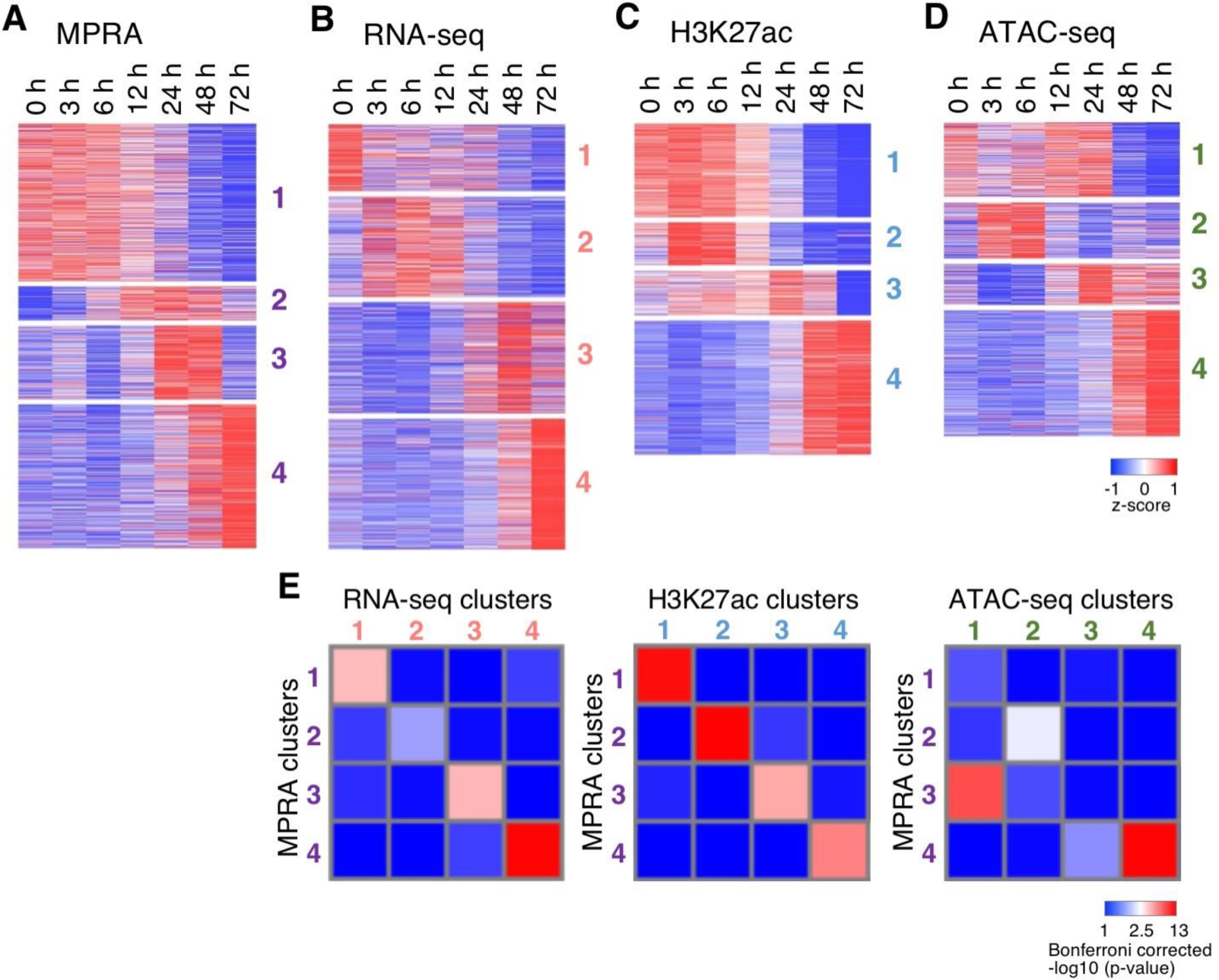
Activity of temporal enhancers: comparing lentiMPRA to the endogenous signals. (**A-D**) Temporal signal of MPRA activity (A), the corresponding signal from an overlapping peak of ATAC-seq **(B)**, H3K27ac ChIP-seq **(C)** and the closest gene signal detected by RNA-seq **(D)** clustered into four temporal groups separately. Values shown are RNA/DNA ration (for for lentiMPRA) and normalized read counts (for all others). Rows are standardized. **(E)** Overlap between the lentiMPRA clusters and the three genomic data modalities. Shown are Bonferroni corrected p-values of a hypergeometric test. The overlap is computed either at the region level (lentiMPRA vs. ATAC-seq or ChIP-seq) or at the gene level (lentiMPRA vs. RNA-seq; using the nearest gene to represent each lentiMPRA region).

Starting with the expression of the closest gene, we find significant levels of overlap between the respective clusters (lentiMPRA and RNA-seq; Bonferroni corrected hypergeometric p-value<0.05; **Fig. 4E**). The significant overlap is observed primarily in time-matched clusters, indicating that an overall trend in the data is that the temporal enhancers are capable of inducing reporter gene expression that is similar to the (postulated) endogenous target gene. Indeed, in an alternative analysis, we defined the *maximal segment* of each enhancer as the two subsequent time points in which it reaches its maximal expression. Comparing the MPRA and the endogenous mRNA, we found that in 48% (752/1,547) of the temporal enhancers the respective maximal segments overlap. Gene ontology enrichment analyses for the genes associated by proximity with the regions in the different clusters, also found gene categories fitting with the temporal expression (**table S5**). For instance, the early cluster is enriched for recruitment of histone acetyltransferases (HATs) and expression of pluripotent genes (e.g. *KLF4*), indicative of stem cell differentiation processes that take place during these early time points (*19*). The mid-early and mid-late clusters are enriched with early chromatin response and genes that are involved in developmental processes. The late cluster is enriched for open chromatin, HAT recruitment and expression of neural genes (e.g. *OTX1*). We observe similar results for the more restricted set of regions that are temporal and active in at least one time point (**fig. S11**).

We next set out to compare the temporal patterns observed with MPRA to those observed at the chromatin level. As expected, we find that the temporal enhancer regions rarely overlap with H3K27me3 peaks (**fig. S1C**). Conversely, using the concept of maximal segments as above, we find that 50% of the temporal enhancers reach their maximum level around the same time as the respective H3K27ac peak (i.e., 604 out of 1,208 temporal enhancers that intersect with a H3K27ac peak). We also observe a similar level of agreement when comparing lentiMPRA to the temporal profiles of chromatin accessibility (48% or 553 out of 1,140 temporal enhancers that intersect with an ATAC-seq peak). Overall, we observe a substantial level of agreement between MPRA and the endogenous transcriptional changes, with 67% of the temporal enhances (1,038/1,547) consistent with the temporal patters of least one of the endogenous signals (H3K27ac, accessibility, or mRNA expression). These results suggest that the signal captured by lentiMPRA could be relevant for neural induction and that the activity of the endogenous counterparts of the temporal enhancers may be functional during this process.

The overlaps that we observed, however, are not perfect and the temporal profiles of many other MPRA regions are in fact different from their endogenous counterparts. While this may result from inaccuracies in the various assays, it may also point to a biologically-driven cause. To investigate this phenomenon at the chromatin level, we turned to the cluster-level analysis (**Fig. 4E** and **fig. S12**). This analysis was designed to identify cases where regions that exhibit a certain temporal pattern with MPRA are likely to exhibit another pattern in their accessibility or H3K27 acetylation (adjusted p-value<0.05). As a general trend, the results indicate that `inconsistent’ temporal regions tend to become induced (when assayed by MPRA) after the occurrence of chromatin changes in their respective endogenous loci. For instance, we observe a significant overlap between the set of regions that become induced after 24 hours when examined by MPRA (MPRA cluster 3; **Fig. 4A**), and the set of regions that become (or remain) accessible during the preceding time points (ATAC-seq cluster 1; **Fig. 4D**). Furthermore, this pattern of delay is observed more often with chromatin accessibility, compared with H3K27ac. These results could potentially be explained by our previous observations that DNA accessibility precedes H3K27ac during neural induction, which is followed by gene expression changes (**Fig. 1E**), and that the temporal H3K27ac signal is a stronger indicator for MPRA enhancer activity (**Fig. 3A**).

In addition to inconsistency with the chromatin readouts, we also observe temporal enhancers that were active several time points before their postulated target genes (**fig. S12B**) and the opposite, where genes were active before the enhancer (**fig. S12C**). The pattern of MPRA induction before the endogenous mRNA can be rationalized by additional constraints that may exist in the endogenous regions, but not necessarily in the (random) integration sites such as dependence on a wider chromatin context, which may be required to enable transcription. Conversely, the latter pattern (mRNA before MPRA) is harder to rationalize and is more likely a result of the assay’s inaccuracy. To further investigate this, we first excluded cases where there was another nearby gene (looking at the closest four) that was more correlated with the MPRA signal but showed inconsistency with the closest gene mRNA signal (**Methods**), thus accounting for possible enhancer-gene miss-assignment. Counting the number of occurrences of each of the two patterns, we find that the second one (mRNA before MPRA) is of a substantially lower abundance (137 vs. 358 enhancers), and that it is in fact at the level of overlap between random sets (**Fig. 4E**).

## TF binding site analyses identifies important neural induction genes

As the RNA product of MPRA is non-endogenous, it provides an effective way for estimating the effects of TFs on transcription, eliminating the need to account for indirect effects. We utilized this property to pinpoint which TFs could be driving neural induction at the different time points. To this end, we used experimental data from the public domain along with DNA binding motifs to determine the potential binding landscape of a large cohort of TFs across our tested regions. More specifically, we recorded, for each temporal enhancer: 1) its predicted binding sites using Fimo (*31*) with two sets of TF motifs (*32, 37*) and 2) its overlap with TF ChIP-seq peaks in hESCs (*20*) or in hESC-derived ectoderm (120 hours post neural induction using inhibitors of TGFβ, WNT and BMP; (*19*)). The result of this analysis is a binary binding matrix of TFs by regions with entries indicating either potential binding using FDR<10^-4^ for TF motifs or overlap with TF ChIP-seq peaks.

We next employed a strict motif enrichment analysis based on comparing the number of motif hits in regions within each temporal MPRA cluster versus the restricted set of all regions in the MPRA design (FDR<0.05, Hypergeometric test; **table S6**; **Methods**). This analysis was designed to nominate candidate TFs whose activity is specific to certain phases of the differentiation process. Accordingly, we found that motifs of pluripotent factors (e.g. NANOG, POU5F1, SOX2 (*38*)), were enriched in the early cluster. Furthermore, immediate early response factors (ATF, JUN, FOS), which are known to be induced by stimuli and stress (*39*), were enriched in mid-early enhancers. These observations suggested that early-and mid-early clusters may respond to TFs that function in pluripotency maintenance and the cell’s acute response, such as apoptosis, respectively. We also found that both mid-late and late clusters were enriched for cell fate commitment and specification factor binding. Specifically, SOX, OTX, and Class III POU factor motifs were enriched in both mid-late and late enhancers, suggesting that enhancers in these group were the direct targets of these key neural factors.

## Activity score identifies novel TFs that are important for neural induction

To narrow down the list of candidate TFs for a follow up investigation of their effect on neural induction, we defined a *TF activity score,* which represents the potential to affect transcription at each time point. We considered two factors that can influence TF activity at each time point: 1) More than expected amount of active enhancers at that time point that are predicted to be bound by that TF (*40*), suggesting that this TFs may provide a parsimonious explanation for the MPRA signal (*41*); and 2) Induction of the mRNA that codes for the TF, which may also suggest functional importance (*42-44*). For the former, we focused our attention to enhancers in which the temporal MPRA pattern significantly overlap with the endogenous pattern, namely - the sub-clusters of regions pertaining to significant entries in **Fig. 4E**. Each of these `consistent’ sub-cluster represents a different mode of temporal relationship between MPRA and the endogenous genome - e.g., early induction with matched timing of mRNA expression or H3K27 acetylation, or late induction that appears after the establishment of chromatin accessibility. While other active MPRA regions in our data can be of additional interest, we postulate that focusing on temporal regions that are consistent with the major patterns of overlap with the endogenous processes is desirable when integrating additional genomic readouts (TF binding), and may also increase the odds that the respective endogenous region is indeed functional.

To compute the activity score of each TF (represented by a motif or a ChIP-seq experiment) at each time point, we look for consistent sub-clusters that peak during that time point (in terms of MPRA signal) and that significantly overlap with the putative target regions of the TF (p-value<0.005, Hypergeometric test). We then count the number of putative target regions that appear in at least one significantly overlapping sub-cluster. The final score is defined by the number of regions found at each time point divided by the total number of regions found across all time points. As an additional constraint, we only consider time points in which the mRNA that encodes for the TF is highly expressed (6th or higher quantile of expressed genes) and significantly induced compared to the preceding time point [p-value <10^-5^; (*24*); for the first time point (0 hour), we compare to the subsequent time point (3 hours)].

The resulting TF activity matrix (**Fig. 5A; table S7**) provided a catalog of 107 TFs that could potentially function as regulators of neural induction. Repeating this analysis with the stricter set of temporal regions that were also detected by the per-time point analysis yielded reproducible results (94 out of 107 cataloged TFs were detected). Similar to previous analyses, we clustered the TF activity score to four representative patterns of activity: early, mid-early, mid-late and late response. Overall, we observed an agreement between known hESC and neural induction associated TFs and their temporal time points. For example, in the early cluster, the pluripotent marker NANOG showed high TF activity score at 0 hour, and immediate-early gene products, ATF3, MYC and EGR1, which likely mediate acute response, showed high score at 3 hours, as expected (*39*). TFs that had a high score at later time points (24-72 hours) included several neural TFs, such as SOX1, OTX2 and PAX6.

**Fig. 5.**
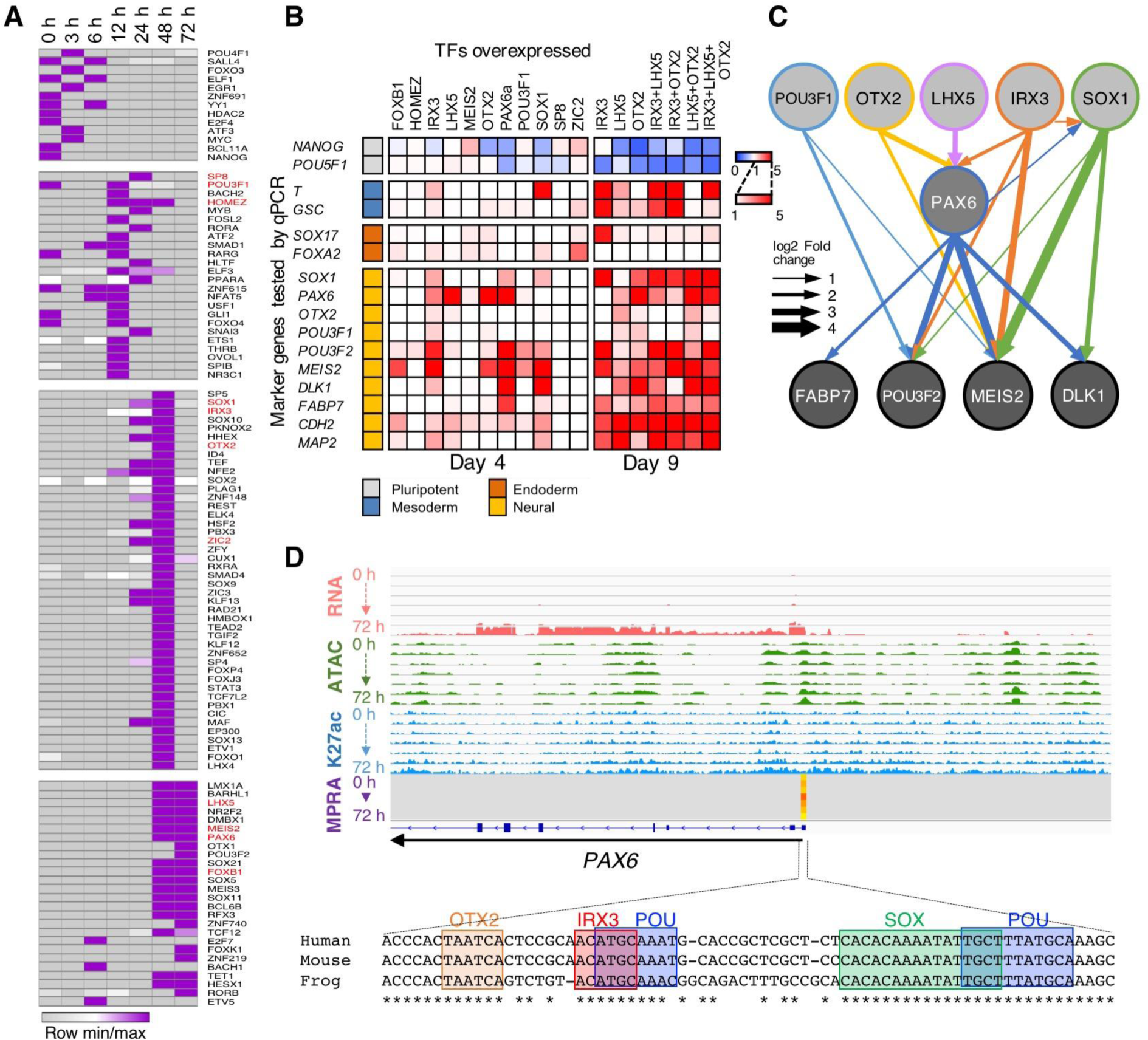
Activity score identifies novel TFs involved in neural induction. (**A**) Heatmap of activity scores per TF per time point. Values are normalized (minimum to maximum) per each row. The eleven TFs used for overexpression (FOXB1, HOMEZ, IRX3, LHX5, MEIS2, OTX2, PAX6, POU3F1, SOX1, SP8 and ZIC2) are marked in red font. (**B**) TF overexpression. The heatmap shows the relative expression of marker genes (pluripotent, mesoderm, endoderm and neural) compared to the *HPRT* gene as determined by qRT-PCR. Results are shown as fold change compared to EGFP overexpression. (**C**) TF overexpression results at day 4 shown as a network. The thickness of arrows represents the log2 fold change. (**D**) TF analyses of the *PAX6* promoter region shows binding sites for OTX2, IRX3, POU and SOX that are evolutionally conserved between human, mouse and frog (*Xenopus tropicalis*).

## Overexpression identifies novel neural induction associated TFs

To test whether our identified TFs are indeed involved in neural induction, we selected for follow up studies eleven representative TFs that were predicted as active during different time points of the induction process: FOXB1, HOMEZ, IRX3, HLX5, MEIS2, OTX2, PAX6, POU3F1, SOX1, SP8, and ZIC2. All of these TFs were selected because their mouse orthologs were shown to be expressed in the neuroectoderm via whole-mount *in situ* hybridizations at E5.5-E8.5, when neural differentiation takes place (*45-55*). Overexpression of PAX6a (short isoform of PAX6) is known to function as a neuroectoderm fate determinant and was previously shown to induce hESCs into a neural lineage (*56*) and was thus selected as a positive control.

We tested if overexpression of each selected TF is sufficient to induce neural induction. The TFs were individually cloned into lentivirus mammalian expression vectors and infected into hESCs. Four days post infection, cells were harvested and RT-qPCR was carried out for lineage marker genes to assess neural induction. The effects on the marker gene expression by each overexpressed TF were normalized to those by EGFP, which was used as a negative control as it is not expected to affect gene expression. Results were represented as a heatmap matrix (**Fig. 5B**) and significant expression changes (comparing to EGFP overexpression using ks-test across the 3 replicates; p-value<0.05) were shown as a network (**Fig. 5C**). We found that overexpression of OTX2, LHX5, and IRX3 were sufficient to induce *PAX6* expression four days after vector transduction, suggesting that these TFs play a role in neural fate specification. IRX3 overexpression also induced other neural markers, such as *SOX1*, *POU3F2* and *MEIS2* directly or indirectly via PAX6 (**Fig. 5C**). However, overexpression of SOX1 or MEIS2 by themselves did not induce *PAX6* expression. SOX1 did however lead to the upregulation of many other neural genes.

Previous studies have shown that *OTX2* overexpression promotes *PAX6* expression in hESCs upon treatment with the TGFβ inhibitor SB431542 and FGF2 (*57*). It was also reported that *LHX2*, a paralog of *LHX5*, promotes *PAX6* expression and neural differentiation in hESCs (*58*). However, despite *LHX5* being expressed in NPCs, its overexpression has yet to be associated with neural induction. The same holds true for *IRX3*, which is known to be expressed in the anterior neuroectoderm in the mouse embryo (*51*), but whose function in neural induction has not been evaluated. Consistent with these findings, analysis of the *PAX6* promoter region found binding sites of OTX, IRX3, SOX and POU that are conserved among species, and were shown by our lentiMPRA to be active around 12-24 hours (**Fig. 5D**), when these TFs are significantly expressed (**Fig. 2A**) and start to gain a high TF activity score (**Fig. 5A**). In addition, we also found several additional examples of functional neural enhancers that contain conserved OTX, SOX, IRX, and/or homeo-domain binding motifs upstream from the *LHX5*, *POU3F2* and *OTX2* genes (**fig. S13**).

As OTX2, LHX5, and IRX3 were able to induce *PAX6* expression and their postulated binding sites are present in many neural induction relevant promoters and enhancers, we tested whether they alone or in combination could lead to a more established neural lineage. This was done by testing the expression of late neural marker genes (i.e. *DLK1*, *FABP7*, *CDH2*, and *MAP2*) nine days after infection of these TFs. We observed upregulation of the late neural markers at day nine (**Fig. 5B**), consistent with the observation that these three factors activated the neural lineage determinant *PAX6* at an early stage. In terms of individual TFs, *LHX5* and *OTX2* upregulated the majority of neural markers we assayed. However, we also observed that overexpression of all three TFs individually or most combinations promoted mesoderm (*T* and *GSC*) and endoderm (*SOX17* and *FOXA2*) markers at day nine. Interestingly, the combination of *LXH5* and *OTX2* together leads to a reduction in expression of these non-neural markers along with a strong neural fate (**Fig. 5B**). In summary, these results suggest that *OTX2, LHX5*, and *IRX3* could function as key regulators of neural differentiation at least partially by activating *PAX6* expression and that neural fate commitment is fine-tuned by combinations of regulators and suppressors that combined lead to this trajectory.

## Discussion

Induced *in vitro* differentiation of embryonic stem cells into neuronal cells is a widely used tool for studying neuronal disease subtypes and for the general understanding of neural development. However, we still do not have a systematic understanding of the genes and regulatory elements that control the initial stages (induction) of this process. Here, we undertook a comprehensive genomic approach to address this and identify molecular components that control gene expression changes during neural induction. We identified previously characterized players in this process along with numerous novel ones. We functionally characterized over two thousand sequences for enhancer activity, finding many to be temporal enhancers. Using all of our genomic data, we put together a combined TF activity score, allowing the identification and experimental validation of novel TFs that are involved in neural induction.

Genomic analyses of multiple time points during early neural induction provided several important findings. We observed that neural induction first involves the silencing of pluripotent markers and upregulation of immediate early genes, corresponding to the cell’s stress response against differentiation stimuli. This is then followed by the upregulation of genes involved in neural lineage fate specification. We also observed that this process is first controlled by chromatin accessibility or simultaneously with H3K27ac modification followed by an increase in mRNA expression. These results also support previous reports about the importance of H3K27ac as an active enhancer mark that correlates with (and possibly affects) temporal changes in transcription levels, which are not captured by accessibility alone (*59*). Finally, our work provides an important catalog of dynamically changing genes and regulatory elements during neural induction.

The use of lentiMPRA allowed us to functionally test thousands of sequences for enhancer activity. A large proportion (68%) of the sequences we tested functioned as active enhancers and 63% showed temporal enhancer activity. We observed a strong temporal correlation between functional enhancers and H3K27ac, but less so with ATAC-seq, likely due to chromatin accessibility preceding enhancer activation. While we observed an overall strong correlation between temporal enhancers and gene expression, it is important to note that this overlap was not obtained for all sequences. We did observe functional enhancers that were also active several time points before their postulated target genes and the opposite, where genes were active before the enhancer. For the latter, our analyses suggest that these discrepancies can stem from various technical factors of the assay. These could include the length of the assayed sequence (171 bp), testing sequences outside their genomic context and other factors.

Analysis of temporal genes and enhancers is important not only to understand the regulatory network underling neural induction, but also to dissect neurological disease. A large body of evidence suggests that the temporal alteration of genes and regulatory elements involved in neuronal development can lead to neurological disease. For example, previous findings have shown that genetic variation influencing cognition and brain size act during neurogenesis (*60*). Fitting with these studies, we observed significant overlap between regions with induced H3K27ac histone modification and neurological disorder GWAS variants. We also observed a significant correlation between these peaks and eQTL variants in general and specifically from brain tissues. Since the statistical background we used for the analysis was the entire set of H3K27ac peaks (regardless of how they change over time), these results suggest that the temporal aspect adds important information, which allowed us to highlight phenotypically important regions.

Our large genomic assays enabled the creation of a TF activity matrix that ranks TF’s temporal functionalities. We identified *OTX2*, *LHX5* and *IRX3* as key regulators of neural induction, as overexpression of these factors were sufficient to induce *PAX6* and other neural markers. While *PAX6* expression in hESCs was shown to be upregulated via *OTX2* (*57*), this finding was novel for *LHX5* and *IRX3*. While *LHX5* is a commonly used neural marker, its ability to induce neural induction was not tested. *IRX3*, was of particular interest. In our study, we found its expression to increase at 12-72 hours, and it obtained a high activity score at these time points, suggesting an important role in neural induction. In our overexpression experiments, we observed that it could by itself induce several neural markers in the hESC culture condition, including *PAX6*. To our knowledge, this is the first report demonstrating a potential role for *IRX3* in neural induction. Although we identified key regulators of neural induction, we also observed that overexpression of different TFs induced different sets of neural markers or even non-neural markers. This observation suggests that the orchestration of multiple TFs is necessary to fine tune neural differentiation. Assays that target the molecular function or regulatory grammar of these key regulators will be necessary in order to further understand this regulatory network.

## Acknowledgments

We want to thank Encoded Therapeutics for assistance on the RNA-seq studies and David Fischer, Martin Kircher, Beth Martin and Jay Shendure (UW) for assistance with the MPRA studies and analyses

## Author contributions

F.I., A.K., N.A. and N.Y. conceived and designed the study. F.I. performed the RNA-seq, ChIP-seq, ATAC-seq, lentiMPRA and overexpression experiments. A.K. and N.Y. preformed the computational analyses, T.A. wrote MPRAanlyze and F.I., A.K., N.A. and N.Y. analyzed the data. N.A. and N.Y. provided resources and F.I., A.K., N.A. and N.Y. wrote the manuscript

## Data and materials availability

All sequencing data is available in the NCBI Gene Expression Omnibus (GEO) as accession number GSE115046.

## Supplementary Materials

Materials and Methods

Supplementary Text

Figures S1-S13

Tables S1-S8

Note S1

File S1

## References and Notes

1. I. Munoz-Sanjuan, A. H. Brivanlou, Neural induction, the default model and embryonic stem cells. Nature reviews. Neuroscience 3, 271–280 (2002).

2. S. M. Chambers et al., Highly efficient neural conversion of human ES and iPS cells by dual inhibition of SMAD signaling. Nat. Biotechnol. 27, 275–280 (2009).

3. N. Yosef, A. Regev, Writ large: Genomic dissection of the effect of cellular environment on immune response. Science 354, 64–68 (2016).

4. F. Inoue, D. Kurokawa, M. Takahashi, S. Aizawa, Gbx2 directly restricts Otx2 expression to forebrain and midbrain, competing with class III POU factors. Molecular and cellular biology 32, 2618–2627 (2012).

5. D. Kurokawa et al., Regulation of Otx2 expression and its functions in mouse forebrain and midbrain. Development 131, 3319–3331 (2004).

6. N. Takasaki, D. Kurokawa, R. Nakayama, J. Nakayama, S. Aizawa, Acetylated YY1 regulates Otx2 expression in anterior neuroectoderm at two cis-sites 90 kb apart. The EMBO journal 26, 1649–1659 (2007).

7. D. W. Volk, D. A. Lewis, Prenatal ontogeny as a susceptibility period for cortical GABA neuron disturbances in schizophrenia. Neuroscience 248C, 154–164 (2013).

8. K. E. Samocha et al., A framework for the interpretation of de novo mutation in human disease. Nat Genet 46, 944–950 (2014).

9. H. O. Kalkman, A review of the evidence for the canonical Wnt pathway in autism spectrum disorders. Mol Autism 3, 10 (2012).

10. N. M. Williams et al., Rare chromosomal deletions and duplications in attention-deficit hyperactivity disorder: a genome-wide analysis. Lancet 376, 1401–1408 (2010).

11. D. J. Roybal et al., Biological evidence for a neurodevelopmental model of pediatric bipolar disorder. The Israel journal of psychiatry and related sciences 49, 28–43 (2012).

12. M. T. Maurano et al., Systematic localization of common disease-associated variation in regulatory DNA. Science 337, 1190–1195 (2012).

13. L. A. Hindorff et al., Potential etiologic and functional implications of genome-wide association loci for human diseases and traits. Proceedings of the National Academy of Sciences 106, 9362–9367 (2009).

14. S. J. Sanders et al., Whole genome sequencing in psychiatric disorders: the WGSPD consortium. Nature neuroscience 20, 1661–1668 (2017).

15. R. Andersson et al., An atlas of active enhancers across human cell types and tissues. Nature 507, 455–461 (2014).

16. A. Fort et al., Deep transcriptome profiling of mammalian stem cells supports a regulatory role for retrotransposons in pluripotency maintenance. Nature genetics 46, 558–566 (2014).

17. J. R. Dixon et al., Chromatin architecture reorganization during stem cell differentiation. Nature 518, 331–336 (2015).

18. W. Xie et al., Epigenomic analysis of multilineage differentiation of human embryonic stem cells. Cell 153, 1134–1148 (2013).

19. A. M. Tsankov et al., Transcription factor binding dynamics during human ES cell differentiation. Nature 518, 344–349 (2015).

20. C. A. Gifford et al., Transcriptional and epigenetic dynamics during specification of human embryonic stem cells. Cell 153, 1149–1163 (2013).

21. E. P. Consortium et al., An integrated encyclopedia of DNA elements in the human genome. Nature 489, 57–74 (2012).

22. F. Inoue et al., A systematic comparison reveals substantial differences in chromosomal versus episomal encoding of enhancer activity. Genome Res. 27, 38–52 (2017).

23. J. Sander, J. L. Schultze, N. Yosef, ImpulseDE: detection of differentially expressed genes in time series data using impulse models. Bioinformatics 33, 757–759 (2017).

24. M. I. Love, W. Huber, S. Anders, Moderated estimation of fold change and dispersion for RNA-seq data with DESeq2. Genome Biol. 15, 550 (2014).

25. A. Subramanian et al., Gene set enrichment analysis: a knowledge-based approach for interpreting genome-wide expression profiles. Proc. Natl. Acad. Sci. U. S. A. 102, 15545–15550 (2005).

26. C. Y. McLean et al., GREAT improves functional interpretation of cis-regulatory regions. Nat. Biotechnol. 28, 495–501 (2010).

27. W. Herzog, K. Weber, Fractionation of brain microtubule-associated proteins. Isolation of two different proteins which stimulate tubulin polymerization in vitro. European journal of biochemistry 92, 1–8 (1978).

28. M. Endo, R. Doi, M. Nishita, Y. Minami, Ror family receptor tyrosine kinases regulate the maintenance of neural progenitor cells in the developing neocortex. Journal of cell science 125, 2017–2029 (2012).

29. G. T. Consortium, Human genomics. The Genotype-Tissue Expression (GTEx) pilot analysis: multitissue gene regulation in humans. Science 348, 648–660 (2015).

30. R. Leslie, C. J. O’Donnell, A. D. Johnson, GRASP: analysis of genotype-phenotype results from 1390 genome-wide association studies and corresponding open access database. Bioinformatics 30, i185–194 (2014).

31. C. E. Grant, T. L. Bailey, W. S. Noble, FIMO: scanning for occurrences of a given motif. Bioinformatics 27, 1017–1018 (2011).

32. P. Kheradpour, M. Kellis, Systematic discovery and characterization of regulatory motifs in ENCODE TF binding experiments. Nucleic Acids Res. 42, 2976–2987 (2014).

33. A. Kreimer et al., Predicting gene expression in massively parallel reporter assays: A comparative study. Human mutation, (2017).

34. A. Kreimer, Y. Zhongxia, N. Ahituv, N. Yosef, Meta-analysis of massive parallel reporter assay enables functional regulatory elements prediction. BioRxiv, (2017).

35. Q. Wu et al., Sall4 interacts with Nanog and co-occupies Nanog genomic sites in embryonic stem cells. The Journal of biological chemistry 281, 24090–24094 (2006).

36. D. J. Rodda et al., Transcriptional regulation of nanog by OCT4 and SOX2. The Journal of biological chemistry 280, 24731–24737 (2005).

37. M. T. Weirauch et al., Determination and inference of eukaryotic transcription factor sequence specificity. Cell 158, 1431–1443 (2014).

38. L. A. Boyer et al., Core transcriptional regulatory circuitry in human embryonic stem cells. Cell 122, 947–956 (2005).

39. H. R. Herschman, Primary response genes induced by growth factors and tumor promoters. Annual review of biochemistry 60, 281–319 (1991).

40. S. R. Grossman et al., Systematic dissection of genomic features determining transcription factor binding and enhancer function. Proceedings of the National Academy of Sciences of the United States of America 114, E1291–E1300 (2017).

41. N. Kashtan, U. Alon, Spontaneous evolution of modularity and network motifs. Proceedings of the National Academy of Sciences of the United States of America 102, 13773–13778 (2005).

42. Y. Setty, A. E. Mayo, M. G. Surette, U. Alon, Detailed map of a cis-regulatory input function. Proceedings of the National Academy of Sciences of the United States of America 100, 7702–7707 (2003).

43. N. Rosenfeld, J. W. Young, U. Alon, P. S. Swain, M. B. Elowitz, Gene regulation at the single-cell level. Science 307, 1962–1965 (2005).

44. N. Yosef et al., Dynamic regulatory network controlling TH17 cell differentiation. Nature 496, 461–468 (2013).

45. S. L. Ang et al., The formation and maintenance of the definitive endoderm lineage in the mouse: involvement of HNF3/forkhead proteins. Development 119, 1301–1315 (1993).

46. T. Inoue, S. Nakamura, N. Osumi, Fate mapping of the mouse prosencephalic neural plate. Developmental biology 219, 373–383 (2000).

47. M. Oulad-Abdelghani et al., Meis2, a novel mouse Pbx-related homeobox gene induced by retinoic acid during differentiation of P19 embryonal carcinoma cells. Developmental dynamics: an official publication of the American Association of Anatomists 210, 173–183 (1997).

48. H. Z. Sheng et al., Expression of murine Lhx5 suggests a role in specifying the forebrain. Developmental dynamics: an official publication of the American Association of Anatomists 208, 266–277 (1997).

49. P. Elms et al., Overlapping and distinct expression domains of Zic2 and Zic3 during mouse gastrulation. Gene expression patterns: GEP 4, 505–511 (2004).

50. L. H. Pevny, S. Sockanathan, M. Placzek, R. Lovell-Badge, A role for SOX1 in neural determination. Development 125, 1967–1978 (1998).

51. A. Bosse et al., Identification of the vertebrate Iroquois homeobox gene family with overlapping expression during early development of the nervous system. Mechanisms of development 69, 169–181 (1997).

52. A. Simeone et al., A vertebrate gene related to orthodenticle contains a homeodomain of the bicoid class and demarcates anterior neuroectoderm in the gastrulating mouse embryo. The EMBO journal 12, 2735–2747 (1993).

53. D. Bayarsaihan et al., Homez, a homeobox leucine zipper gene specific to the vertebrate lineage. Proceedings of the National Academy of Sciences of the United States of America 100, 10358–10363 (2003).

54. Q. Zhu et al., The transcription factor Pou3f1 promotes neural fate commitment via activation of neural lineage genes and inhibition of external signaling pathways. eLife 3, (2014).

55. W. C. Dunty, Jr., M. W. Kennedy, R. B. Chalamalasetty, K. Campbell, T. P. Yamaguchi, Transcriptional profiling of Wnt3a mutants identifies Sp transcription factors as essential effectors of the Wnt/beta-catenin pathway in neuromesodermal stem cells. PloS one 9, e87018 (2014).

56. X. Zhang et al., Pax6 is a human neuroectoderm cell fate determinant. Cell stem cell 7, 90–100 (2010).

57. B. Greber et al., FGF signalling inhibits neural induction in human embryonic stem cells. The EMBO journal 30, 4874–4884 (2011).

58. P. S. Hou et al., LHX2 regulates the neural differentiation of human embryonic stem cells via transcriptional modulation of PAX6 and CER1. Nucleic acids research 41, 7753–7770 (2013).

59. N. D. Heintzman et al., Histone modifications at human enhancers reflect global cell-type-specific gene expression. Nature 459, 108–112 (2009).

60. L. de la Torre-Ubieta et al., The Dynamic Landscape of Open Chromatin during Human Cortical Neurogenesis. Cell 172, 289–304 e218 (2018).

61. J. D. Buenrostro, P. G. Giresi, L. C. Zaba, H. Y. Chang, W. J. Greenleaf, Transposition of native chromatin for fast and sensitive epigenomic profiling of open chromatin, DNA-binding proteins and nucleosome position. Nat. Methods 10, 1213–1218 (2013).

62. D. Kim et al., TopHat2: accurate alignment of transcriptomes in the presence of insertions, deletions and gene fusions. Genome Biol. 14, R36 (2013).

63. A. M. Bolger, M. Lohse, B. Usadel, Trimmomatic: a flexible trimmer for Illumina sequence data. Bioinformatics 30, 2114–2120 (2014).

64. C. Trapnell et al., Transcript assembly and quantification by RNA-Seq reveals unannotated transcripts and isoform switching during cell differentiation. Nat. Biotechnol. 28, 511–515 (2010).

65. Y. Liao, G. K. Smyth, W. Shi, featureCounts: an efficient general purpose program for assigning sequence reads to genomic features. Bioinformatics 30, 923–930 (2014).

66. B. Langmead, C. Trapnell, M. Pop, S. L. Salzberg, Ultrafast and memory-efficient alignment of short DNA sequences to the human genome. Genome Biol 10, R25 (2009).

67. B. Langmead, S. L. Salzberg, Fast gapped-read alignment with Bowtie 2. Nat Methods 9, 357–359 (2012).

68. Y. Zhang et al., Model-based analysis of ChIP-Seq (MACS). Genome Biol. 9, R137 (2008).

69. A. R. Quinlan, I. M. Hall, BEDTools: a flexible suite of utilities for comparing genomic features. Bioinformatics 26, 841–842 (2010).

70. A. R. Quinlan, I. M. Hall, BEDTools: a flexible suite of utilities for comparing genomic features. Bioinformatics 26, 841–842 (2010).

71. J. MacArthur et al., The new NHGRI-EBI Catalog of published genome-wide association studies (GWAS Catalog). Nucleic Acids Res 45, D896–d901 (2017).

72. J. Malone et al., Modeling sample variables with an Experimental Factor Ontology. Bioinformatics 26, 1112–1118 (2010).

73. A. D. Johnson et al., SNAP: a web-based tool for identification and annotation of proxy SNPs using HapMap. Bioinformatics 24, 2938–2939 (2008).

74. W. J. Kent et al., The human genome browser at UCSC. Genome research 12, 996–1006 (2002).

75. F. Inoue et al., A systematic comparison reveals substantial differences in chromosomal versus episomal encoding of enhancer activity. Genome Res. 27, 38–52. doi: 10.1101/gr.212092.212116. Epub 212016 Nov 212099. (2017).

76. J. Zhang, K. Kobert, T. Flouri, A. Stamatakis, PEAR: a fast and accurate Illumina Paired-End reAd mergeR. Bioinformatics 30, 614–620 (2013).

77. H. Li, R. Durbin, Fast and accurate short read alignment with Burrows-Wheeler transform. Bioinformatics 25, 1754–1760 (2009).

